# PyInteraph2 and PyInKnife2 to analyze networks in protein structural ensembles

**DOI:** 10.1101/2020.11.22.381616

**Authors:** Valentina Sora, Matteo Tiberti, Shahriyar Mahdi Robbani, Joshua Rubin, Elena Papaleo

## Abstract

**Motivation:** Protein dynamic is essential for cellular functions. Due to the complex nature of non-covalent interactions and their long-range effects, the analysis of protein conformations using network theory can be enlightening. Protein Structure Networks (PSNs) rely on different philosophies, and the currently available tools suffer from limitations in terms of input formats, supported network models, and version control. Another issue is the precise definition of cutoffs for the network calculations and the assessment of the stability of the parameters, which ultimately affect the outcome of the analyses.

**Results:** We provide two open-source software packages, i.e., PyInteraph2 and PyInKnife2, to implement and analyze PSNs in a harmonized, reproducible, and documented manner. PyInteraph2 interfaces with multiple formats for protein ensembles and calculates a diverse range of network models with the possibility to integrate them into a macro-network and perform further downstream graph analyses. PyInKnife2 is a standalone package that supports the network models implemented in PyInteraph2. It employs a jackknife resampling approach to estimate the convergence of network properties and streamline the selection of distance cutoffs. Several functionalities are based on MDAnalysis and NetworkX, including parallelization, and are available for Python 3.7. PyInteraph2 underwent a massive restructuring in terms of setup, installation, and test support compared to the original PyInteraph software.

**Conclusions:** We foresee that the modular structure of the code and the version control system of GitHub will promote the transition to a community-driven effort, boost reproducibility, and establish harmonized protocols in the PSN field. As developers, we will guarantee the introduction of new functionalities, assistance, training of new contributors, and maintenance of the package.

**Availability:** The packages are available at https://github.com/ELELAB/pyinteraph2 and https://github.com/ELELAB/PyInKnife2 with guides provided within the packages.

## Introduction

Proteins are highly dynamic entities, and even the knowledge of sparsely populated conformational states of proteins is key to understanding their function (Baldwin and Kay, 2009).

The structural ensemble that a protein can attain in its folded state entails a rugged free-energy landscape, where the main free energy minimum corresponds to the major state of the protein, and other local minima constitute alternative ‘minor’ states (Motlagh *et al*., 2014). Perturbations such as a binding event with another biomolecule, a mutation, or a post-translational modification, may affect the population of these states. For example, perturbations can switch a minor state to a more frequently occurring state (Naganathan, 2019; Guarnera and Berezovsky, 2019; Abrusán and Marsh, 2019; Papaleo, Sutto, *et al*., 2014; Lambrughi, De Gioia, *et al*., 2016). Distal residues can influence the transitions between different states, mechanisms which are at the base of allostery (del Sol *et al*., 2009; Tsai and Nussinov, 2014; Papaleo, 2015; Papaleo *et al*., 2016; Ribeiro and Ortiz, 2016).

In recent years, we witnessed enormous progress in experimental and computational structural biology to study protein dynamics and allostery (Papaleo, 2015; Papaleo *et al*., 2016; Lisi and Loria, 2016; van den Bedem and Fraser, 2015; Keedy *et al*., 2015). In this context, due to the complex nature of intra- and inter-molecular weak interactions and their capability to exert effects over long distances in the structure, the description of long-range communication and protein structures using network theory provides an asset (Di Paola *et al*., 2013; Di Paola and Giuliani, 2015; Vuillon and Lesieur, 2015).

In this context, the non-covalent intra- and intermolecular interactions in proteins are fundamental to determine their three-dimensional structures and the changes among different states. They can be represented as a network, namely a Protein Structure Network (PSN). PSNs are generally small-world networks (Atilgan *et al*., 2004; Vendruscolo *et al*., 2002) making them suitable for the fast transmission of conformational changes among different distal sites. In these small-world networks, the residues communicate through the shortest paths available, and multiple routes of communication are in play and often pass through common nodes (del Sol *et al*., 2009; Invernizzi *et al*., 2014; Meireles *et al*., 2011).

The usage of network-based approaches to protein dynamics is recent and still developing (Papaleo *et al*., 2012; Invernizzi *et al*., 2014; Nygaard *et al*., 2016; Mariani *et al*., 2013; Angelova *et al*., 2011; Ghosh and Vishveshwara, 2007; Astl and Verkhivker, 2019). Proper reproducible protocols and solid tools accessible to the community are needed in the PSN field to reach the standards of recent open-access and collaborative initiatives for reproducibility and transparency in molecular modeling and simulations (PLUMED Consortium, 2019; Smith *et al*., 2020; Senapathi *et al*., 2020). These initiatives are also common in other areas of bioinformatics, such as cancer genomics (Siu *et al*., 2016; Colaprico *et al*., 2020, 2015; Mounir *et al*., 2019; Terkelsen *et al*., 2020).

In 2014, we developed PyInteraph (Tiberti *et al*., 2014)for the study of protein structure networks (PSNs) from structural ensembles, especially suited to work on trajectories from atomistic simulations such as Molecular Dynamics (MD). Examples of PyInteraph applications includes: i) the study of the effects of mutations in disease-related proteins (Nygaard *et al*., 2016; Kønig *et al*., 2019; Marino *et al*., 2015; Pantsar *et al*., 2018; Lambrughi, Lucchini, *et al*., 2016; Kumar and Papaleo, 2020; Fas *et al*., 2019; Endo *et al*., 2020; Michelini *et al*., 2020; Di Stazio *et al*., 2020), ii) to characterize or design variants for enzymes of industrial interest (Jónsdóttir *et al*., 2014; Papaleo, Parravicini, *et al*., 2014; Michetti *et al*., 2017; Óskarsson *et al*., 2016; Singh *et al*., 2016), to study the binding of biomolecules to a target protein (Di Rita *et al*., 2018), and to disclose the effect of post-translational modifications (Faienza *et al*., 2020; Lambrughi, De Gioia, *et al*., 2016). More broadly, PyInteraph has been used to study protein dynamics and allostery (Lambrughi, De Gioia, *et al*., 2016; Sora and Papaleo, 2019; Marino and Dell’Orco, 2016, 2019; Galochkina *et al*., 2019; Faienza *et al*., 2020; Abbas *et al*., 2019; Bonì *et al*., 2020; Borsatto *et al*., 2019). PyInteraph is also included in planned protocols from scientific consortia to study the genetics and the molecular mechanisms at the base of different diseases (Borges *et al*., 2020). Due to the large and increasing number of applications, it becomes thus of paramount importance to maintain and develop further the package.

Here, we provide a new version for Python 3 upon significant restructuring of the code for a more flexible and accessible structure, i.e., PyInteraph2. Moreover, we developed the support tool PyInKnife2 for analyses of convergence of the network properties and cutoff selection.

## Features and Implementation

PyInteraph2 (Figure 1) calculates the most relevant non-bonded interactions between pairs of residues on structural ensembles and translates such information into a network. Those interactions include salt bridges, hydrogen bonds, and contacts between hydrophobic side chains, along with a description based on a knowledge-based potential for the calculation of energy networks (Potapov *et al*., 2009, 2010). Edges in the resulting networks are weighted on the persistence of the interaction they represent, defined as the percentage of conformations in which the interaction is present in the ensemble. This value is used to understand whether two residues or nodes are connected in the network or not. It also allows for unweighted networks where different interactions can be combined. The knowledge-based potential can also be used to weight the network.

**Figure 1.**
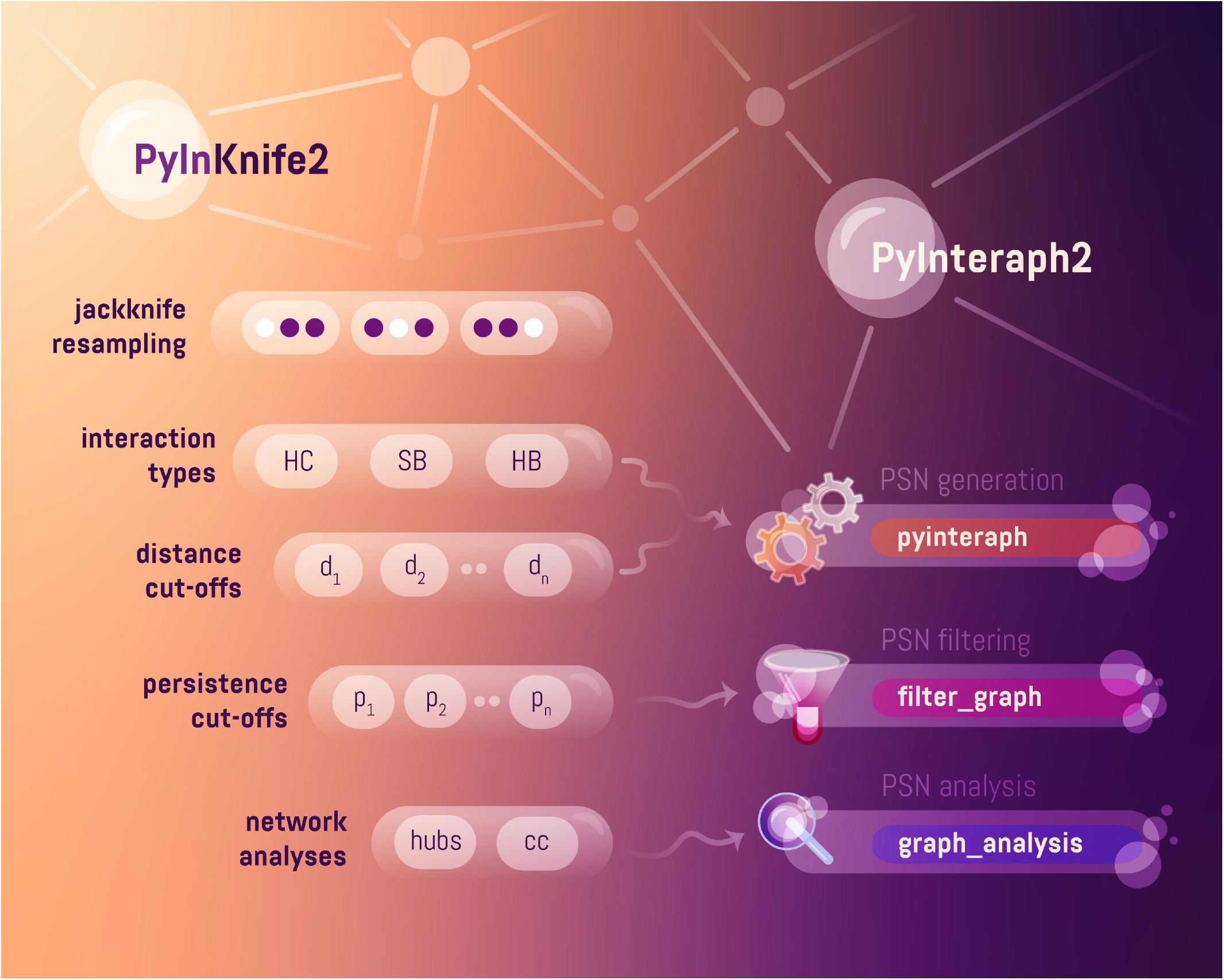
PyInteraph2 and PyInKnife2. Schematic depiction of PyInteraph2 and PyInKnife2 and how PyInKnife2 incorporates PyInteraph2 tools into its workflow. “HC” stands for hydrophobic contacts, “SB” stands for salt bridges, “HB” stands for hydrogen bonds, “cc” stands for connected components.

Graphs for each interaction type are calculated separately, and, once they have been computed, PyInteraph2 allows to process them in several ways. The *pyinteraph* primary function allows the calculation of the network. Also, it provides a table with pairwise intra- and intermolecular interactions and their persistence for further consultation. Usually, a graph is first filtered by removing those edges with low persistence, and the software includes the possibility to identify an optimal persistence threshold (p_crit_). The user can test a range of persistence cutoffs for a given unfiltered PSN. For this analysis, the user can modify the lower limit, upper limit, and step size. Afterward, PyInteraph2 computes the size of the most populated connected component (i.e., the biggest interconnected sub-graph in the main network) for each filtered network. The optimal persistence threshold will be where there is a sharp fall in the number of nodes in the most populated connected component, similar to the definition of the optimal interaction strength (I_crit_) in the PSN based on atomic contacts (Brinda and Vishveshwara, 2005). The user can carry out the p_crit_ analysis and the filtering step with the *filter_graph* function of PyInteraph2. After filtering, it is possible to merge the networks into a single multi-interaction graph.

PyInteraph2 also includes downstream graph analyses that are implemented in the *graph_analysis* function. For example, it allows the identification of hubs, i.e., well-connected nodes in the network, and calculates connected components. PyInteraph2 also enables the calculation of paths between pairs of residues in the network. The analysis of connected components and paths uses NetworkX functions. The functions implement a breadth-first search algorithm to find the connected components and a modified depth-first search algorithm to compute the paths. Hubs are found simply by reporting nodes meeting or exceeding a pre-defined node degree threshold, which one should select after careful reading of the relevant literature. For the most common PSN networks, the threshold used to assign a node as a hub is a degree value of 3 or 4 (Papaleo, 2015).

A path is composed of nodes that can be traversed in the network while trying to reach a target node from a source one. This is particularly useful to identify potential routes of structural communication between distant protein sites - for instance, an active site and an allosteric site.

We also provide the possibility to calculate networks of contacts between residue side chains (scPSN) by taking advantage of the hydrophobic contacts function, which simply considers distances between the centers of mass of residue side chains to identify a contact. Such a network accounts for any type of contact employing a distance cut-off between the pairs of centers of mass as the only required parameter. We benchmarked the method on proteins with different sizes, folds, and simulated using different physical models, to estimate the best distance cut-off to obtain a network with balanced connectivity using a jackknife resampling method (Salamanca Viloria *et al*., 2017).

PyInteraph, which was initially written for Python 2.7, includes C extensions exposed through a Cython abstraction layer. The package uses several open-source packages, such as MDAnalysis (Michaud-Agrawal *et al*., 2011), NetworkX (Hagberg *et al*., 2008), Matplotlib (Hunter, 2007), and the SciPy stack (Harris *et al*., 2020; Virtanen *et al*., 2020) to perform the required operations on a conformational ensemble, as well as to calculate and represent graphs. Its installation required few open-source Python packages, Cython, and a C compiler. Python 2.7 is obsolete, and the support for it has ceased. Most of the commonly used Python packages have transitioned to version 3. Moreover, changes applied over time to the necessary packages had made the installation of PyInteraph complicated, usually requiring an ad-hoc Python virtual environment. We have now ported the PyInteraph2 software to Python 3, having Python 3.7 as the main development target, and discarding retro-compatibility with Python 2. We have updated the source code to be compatible with the latest stable release of requirements at the time of writing, making it trivial to install. Furthermore, we have refactored the structure of the software package to be compliant with the standard formats and features of modern Python packages.

Previously, a full installation of PyInteraph required the user to set up a few system variables to make it work correctly. We exploit features from the *setuptools* package to install the necessary files, Python packages, and scripts at a standard location in the new PyInteraph2 release. As before, the setup script installs two Python packages: i) *pyinteraph*, which includes the most important functions of the package, including user scripts, and ii) *libinteract*, which acts as a backend. Nonetheless, the new structure of user-executable scripts allows to import the functions easily and test them separately. We included more in-depth tests for both packages with automated *pytest* scripts to maintain and expand routinely. The outputs on the *pyinteraph* tool have been modified to better handle ensembles of protein-protein complexes (see as an example: https://github.com/ELELAB/pyinteraph2/tree/master/example).

The current version of PyInteraph2 provides a modular structure of the code, which means that it is possible to add new functionalities without significant alterations of the core part of the code. We also provide guidelines for contributors who are interested in joining the project (https://github.com/ELELAB/pyinteraph2/edit/master/CONTRIBUTE).

We also provide a standalone version of a pipeline for downstream network analysis to identify suitable distance cutoffs for PyInteraph-based PSNs, i.e., PyInKnife2 (Figure 1). PyInKnife2 is also available for Python 3, and it supports all the different networks calculated by PyInteraph2, i.e., hydrogen bonds, salt bridges, hydrophobic interactions, and side-chain-based contact maps. PyInKnife2 has been designed in a user-friendly manner thanks to customizable configuration files instead of several command-line options. This design will facilitate the reproducibility of single runs. Moreover, the calculations are fully parallelized to improve the performances on the analysis of multiple trajectories (ensembles) simultaneously. For this, we decided to rely on the Dask package (Rocklin, 2015) to keep a clean interface and ease scaling the pipeline up or down according to the computational resources available. Furthermore, the customization of the analyses is fine-grained with a modular interface to provide maximum flexibility for the user. Overall, PyInKnife2 provides a unified Python package, consisting of three main executables to run the pipeline, aggregate the results and plot them, and a few modules where the core functions live. PyInteraph2 and PyInKnife2 are available under the GNU General Public Licence (GPL) version 3.0.

## Comparison with other software packages for PSN on protein ensembles

Table S1 contains a comparison of several features of the currently available tools (Bhattacharyya *et al*., 2016; Seeber *et al*., 2011; Serçinolu and Ozbek, 2018; Chakrabarty *et al*., 2019; Brown *et al*., 2017; Grant *et al*., 2020; Contreras-Riquelme *et al*., 2018) to generate PSNs from structural ensembles. We considered: i) features related both to the PSN construction methodology and ii) to the palette of network analyses implemented, together with iii) indicators of the availability of the tools across different platforms and types of user interfaces.

We also assessed the supported formats for the input file containing the structural ensembles and whether a version control repository is available for the source code, where individual users can contribute.

The “tools to support the selection of parameters” field refers to methods provided with the PSN tools or elsewhere, helping the selection of values for optimal cutoffs for one or more network parameters. Only three packages have integrated similar protocols, either internally (PyInteraph2) or referring to external resources (PSN-ensemble and WORDOM, PyInteraph2 with PyInKnife2).

PyInteraph2 provides high flexibility in terms of the supported input formats and the classes of interactions to calculate the network. We noticed that there is a need for the expansion of the supported downstream analyses in PyInteraph beyond the common ones. For example, we will consider the inclusion of important properties such as communities, k-cliques, and centrality measurements provided by other packages in the future.

We discriminated between standalone software packages, web servers, and indicated whether an API for programmatic access was available). In addition, We reported if plug-ins for widely used molecular visualization software packages were available to map the interactions found in the PSNs on the corresponding protein structure to ease the interpretation of the network and subsequent analyses. While NAPS provide some degree of visualization, only PyInterph2 and RIP-MD have a plug-in to integrate the network analyses with software packages for molecular visualization. We noticed that only some projects provide version control repositories. Moreover, in our survey, we found published tools that are no longer maintained. This is a common issue in bioinformatics. The possibility to develop and sustain a centralized resource in a community-driven direction should help to overcome it.

## Case of study

We used the new PyInKnife2 on one microsecond MD trajectory of the Cyclophilin A (CypA) wild-type enzyme, previously published (Papaleo, Sutto, *et al*., 2014; Salamanca Viloria *et al*., 2017). CypA is a well-studied enzyme in terms of allostery and structural communication both with experimental and computational techniques (Ramanathan *et al*., 2011; van den Bedem *et al*., 2013; Rodriguez-bussey *et al*., 2018; Fraser *et al*., 2009; Camilloni *et al*., 2014; McGowan and Hamelberg, 2013; Papaleo, Sutto, *et al*., 2014;

Wapeesittipan *et al*., 2019). PyInKnife2 supports running the full jackknife resampling protocol on the different types of networks implemented in PyInteraph. In this example, we have used PyInKnife2 to run the jackknife approach on the scPSN as an illustrative example (see files in https://github.com/ELELAB/pyinteraph2/tree/master/examples/CypA/graph_analysis/pyinknife).

After setting up our system, by resolving periodic boundary conditions and keeping the protein atoms only in our trajectory, we have prepared the PyInKnife2 YAML format configuration file. This file allows the configuration of all the parameters to perform the jackknife protocol, most notably the number of resamplings. The configuration file also specifies the range of distance cutoffs to use in the jackknife resampling, which is repeated for every point of the selected parameter range to perform a parameter scan. Any other command-line option to be passed to the PyInteraph2 executable scripts can be specified within the configuration file, allowing maximum flexibility. PyInKnife2 uses the configuration file to automatically create the required directory structure, appropriately slice the trajectory, and perform the necessary analyses. The output is a set of neatly organized directories to store the output files of the analysis. A second step allows the aggregation of the data into easily-readable comma-separated files (CSV) containing information about the network properties of interest, i.e., number of hubs and size of connected components. A third script allows visualizing this information as publication-ready plots, requiring a second configuration file to fine-tune the graphical representation of the output.

For the scPSN analysis, we used PyInKnife2 to perform the jackknife resampling with ten windows and to vary the distance parameter in the 4.5-6.0 Å range, every 0.1 Å. We have then considered the size of the five largest connected components and the distribution of the number of hub residues over node degree, where hubs were defined as those residues connected to at least three others. We filtered the networks by removing the edges with weight < 20% to keep the most significant ones.

Our analysis with PyInKnife2 confirms that our investigated network properties are stable with the resampling approach and that 5.0A (figure 2A) is a suitable distance cut-off for this type of network, as already recommended before (Salamanca Viloria *et al*., 2017). Networks at 4.9 Å have very small connected components, as the network is too fragmented. In contrast, networks at higher cut-off clump most residues into a single connected component (as can be seen in https://github.com/ELELAB/pyinteraph2/tree/master/examples/CypA/graph_analysis/pyinknife). PyInKnife2 also has the purpose of assessing the stability of the network parameters calculated from the MD ensemble. Our properties feature low standard errors, suggesting that they are stable within the resampling performed by the jackknife approach. To this goal, we used the results from PyInteraph2 to investigate the scPSN calculated on the full trajectory using the graph_analysis tool of PyInteraph2.

**Figure 2.**
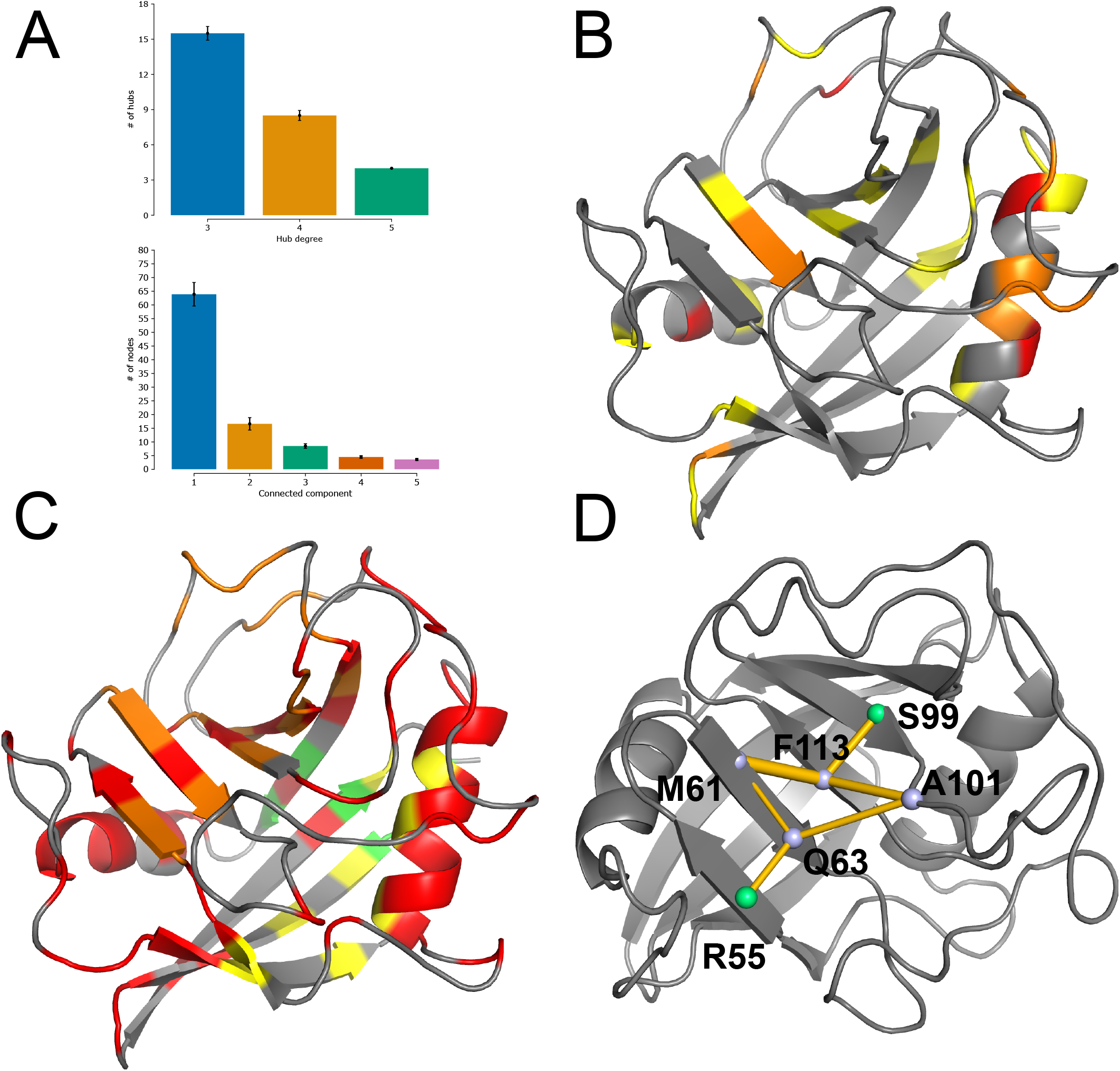
scPSN analysis of CypA. A) Results for hubs and connected components identified in the ensemble using the PyInKnife2 pipeline. B) Hubs identified by PyInteraph2 on the scPSN network. Red, orange, and yellow residues have degrees of 5, 4, and 3, respectively. C) Connected components identified in the scPSN; residues with the same color belong to the same connected component. The largest ones are highlighted in red, orange, yellow, green according to a descending order of size as described in the text. D) Communication paths between S99 and R95.

The scPSN features a total of 30 hub residues (Figure 2B), of which four with degree 5, 10 of degree 4, and 16 of degree 3. They are distributed on the whole structure and present both in stable secondary structure elements and in surface-exposed loops. The α-helix 1 (residues 30-41) is significantly enriched with hub residues, hosting six of them, of which two with the highest degree identified. Hub residues are in all the β-strands of the β-barrel fold of CypA except in the β-strand 3 and loops and turns that connect different secondary structure elements. The highest-degree nodes are T32 and A38, both located on the α-helix 1, H92, on a surface loop and spatially close to S99, and M142, on the α-helix 3. S99 is particularly important since its conformational changes can trigger long-range effects to the catalytic site and its mutation to threonine impair catalytic activity (Fraser *et al*., 2009).

The network of contacts identified by PyInteraph2 consists of several connected components (Figure 2C). The largest connected component accounts for 58 nodes, touches upon most of the protein structure, and spans over the helices α1, α3, and the whole β-sheet. It also includes residues of the loops interconnecting these secondary structure elements. The remaining connected components have a markedly more local character. The second-largest component contains 19 nodes, which are mostly residues in the strands β3-β6. This connected component consists of the long-range communication network described in the next section from the core (S99) to the active site (R55). The third one (11 nodes) has a local character and comprises part of the α1 helix and of the neighboring strands β1, β2, and β8. The fourth component (6 nodes) is also very localized, including residues in the strands β1, β2, and β7. Finally, we have calculated the shortest communication paths in the network between two residues using the *graph_analysis* tool of PyInteraph2 (https://github.com/ELELAB/pyinteraph2/tree/master/examples/CypA/graph_analysis/pyinknife). In detail, we selected one residue of the active site, R55, and a distant residue, S99 (Figure 2D), for which long-range communication was previously studied by resampling the electron density map from X-ray crystallography (Fraser *et al*., 2009). It is unclear if this path entails long-term scale dynamics as suggested by NMR dispersion experiments or a shorter timescale, as suggested by recent MD works (Wapeesittipan *et al*., 2019). Still, it is clear that the mutation of S99 to Thr, which entraps the protein in the minor state, promoting the cascade of collisional clashes with impact on the enzyme activity (Fraser *et al*., 2009). S99 and R55 are the endpoints of a network of residues, passing through F113 and M61, triggering the transition between a major and a minor conformational state of CypA. This mechanism is supported by the resampling of the electron density map of a crystal of the wild-type and S99T CypA variants (Fraser *et al*., 2009). The S99T mutation traps the enzyme in the minor state, significantly reducing the turnover rate. We investigated the network of side-chain contacts that propagate long-range communication using another PSN framework and free energy calculations (Papaleo, Sutto, *et al*., 2014). Interestingly, we found that Q63 is more likely to take part in the transition between major and minor states than M61. In these new analyses, PyInteraph2 identified two communication paths: i) one that includes the residues F113, M61, Q63 and, ii) one with the residues F113, A101, and Q63. The two paths feature similar average persistence (57.4% and 54.1%).

## Conclusions

In the emerging field of PSNs applied to conformational ensembles, we are still far from deriving precise information and predictions from PSNs. Moreover, the field is still suffering from a lack of consensus in the procedure to employ and how to design or implement the network model. Efforts to develop solid platforms and strong foundations to study PSNs of highly dynamic biomolecules are needed. This is especially relevant because of the potential of PSN approaches in complementing experimental studies of important proteins in health and disease. For example, PSNs may help design new protein variants with different stabilities or binding preferences, classify the impact of disease-related variants, or identify druggable allosteric hotspots. Nevertheless, a more organized community, better standards, and a solid framework in the field are still needed to consolidate these applications. Our work in restructuring PyInteraph2 and designing PyInKnife2 is the first pillar toward a community-driven effort in the PSN field applied to biomolecular ensembles to develop harmonized and reproducible protocols, facilitate the maintenance of the tools and the contributions from other researchers. We provided an open-access, easily accessible, and modular structure supported by version control through our GitHub repositories. These features guarantee the baseline for extending their functionalities to include new network models or increase the support for downstream graph analyses. The usage of PyInteraph2 as a common tool for PSN analyses should also facilitate benchmarking efforts and comparisons among different methods. Moreover, pipelines as the ones implemented in PyInKnife2 could guide the selection of optimal distance cutoffs.

We foresee that these are the first steps towards establishing a community-driven effort to contribute to PyInteraph2. We aim to develop PyInteraph2 as a central resource for different PSN methods where the developers, other contributors, and users who apply the packages for their case study can join forces for long-term sustainable and more transparent solutions.

---

**Table S1. Comparison of PyInteraph2 and PyInKnife2 with other software packages for different classes of PSN**. We did not include MDN (Ribeiro and Ortiz, 2015) since the link was unavailable at the time of writing. Also, the RIP-MD web server version did not work at the time of writing, and we decided to refer only to the standalone version in this Table.

